# Signature of random connectivity in the distribution of neuronal tuning curves

**DOI:** 10.1101/2022.08.30.505888

**Authors:** Pierre Orhan, Adrian Duszkiewicz, Adrien Peyrache

**Affiliations:** Montreal Neurological Institute and Hospital, McGill University, Montreal, QC, Canada; Ecole normale supérieure, PSL University, CNRS, Paris, France; Centre for Discovery Brain Sciences, University of Edinburgh, Edinburgh, UK

## Abstract

Understanding the relationship between circuit properties and the organization of neuronal population activity is a fundamental question in neuroscience. The fine tuning of neuronal activity to specific values of environmental or internal features are canonical examples of how information is encoded in the brain, possibly resulting from precisely organized inputs. Yet, in the cortex, finely tuned neurons are often recorded together with neurons whose tuning is much less specific, for example those of inhibitory neurons, and the connectivity statistics accounting for the overall distribution of tuning curves is unclear. Here, using recordings in the mouse head-direction system, we first show both in simulation and analytically that random linear combinations of ideal finely tuned inputs reproduce the distribution of fast-spiking neuron tuning curves, a class of neurons believed to operate in the linear regime. This transformation preserves, on the population level, the singular spectrum of the input tuning curves but the relative power of each singular component is independently distributed in each output cell, leading to a distribution ranging from uni-modal to symmetrically tuned cells. We then generalize the model to a non-linear transformation of the inputs, combined with background inhibition. Using recordings from input neurons in the thalamus, where tuning curves are near-ideal, the model reproduces for various levels of inhibition the entire range of observed neuronal responses in the cortex, from precisely and narrowly tuned neurons to multipeak excitatory cells, as well as symmetrical tuning curves of inhibitory neurons. We replicate these findings in a dataset of hippocampal recordings. In conclusion, the full distribution of tuning curves is a signature of input connectivity statistics, which, for fast-spiking neurons and thalamocortical circuits, is likely to be random rather than specifically organized.

## Introduction

The brain encodes behaviorally-relevant information in neuronal populations and high-level representations emerge from successive transformations of input signals in feed-forward networks. A classic view posits that specific connectivity profiles account for the observed tuning curves, for example in the visual cortex where the response of orientation-selective neurons can be explained by the specific alignment of non-orientation selective presynaptic neurons in the thalamus [1, 2]. Another example is the fly central complex where precise connectivity gives rise to specific neuronal signals for navigation [3]. Yet, these views are challenged by a series of experimental and computational studies arguing against specific connectivity for the emergence of observed neuronal correlates [4, 5, 6, 7, 8]. For example, orientation selectivity in the cortex may result from random inputs from the thalamus [4, 6]. This is also the case in the spatial navigation system, where the activity of head-direction cells [9, 10] and grid cells [11] could emerge from largely random connectivity [12]. However, a clear theoretical understanding of this phenomenon is lacking.

While most simulations focus on the responses of excitatory neurons, the responses of cortical inhibitory cells are also a matter of debate. These neurons, which orchestrate cortical population activity, show tuning to locally encoded features that are usually not uniform but broader than that of excitatory cells [13, 14, 15, 16, 17, 18, 19, 20, 21, 22]. Similarly, the synaptic origin of their tuning is unclear as they are contacted by local excitatory neurons irrespective of their tuning preference [15, 17].

In many cases, the relationship between connectivity and tuning properties is still largely intractable in mammals, begging the question of whether the distribution of tuning curves can provide, at least indirectly, evidence supporting certain types of connectivity profiles. Here, we address this problem by analyzing data from the head-direction (HD) system [23], specifically in the anterodorsal nucleus (ADN) of the thalamus and its main cortical target, the post-subiculum (PoSub). First, we focus on fast-spiking (FS) inhibitory cells which operate in a near-linear regime, as their resting potential is close to their membrane threshold [24]. This property makes it possible to compare their tuning curves to simple linear model of synaptic integration. We show that their tuning is, overall, well accounted for by a random summation of excitatory cells. We derive the distribution of FS cell tuning curves under this model. Next, we generalize this theory to excitatory cells by composing the model with inhibition and non-linear output transformation. We show that introducing a set level of inhibition is sufficient to reproduce all observed tuning curves for both excitatory and inhibitory cells. In addition, we show that the model account for the effect of specific manipulation on cortical cells of thalamic input rate. Last, we apply the model to a dataset of hippocampal recordings; showing that the model can generalize to other features.

## Results

### A simple model of inhibitory cell tuning in the PoSub

In the post-subiculum, FS neurons show tuning curves that are organized following a simple principle: they share, on average, the same Fourier spectrum as excitatory cells [23]. However, while canonical HD cells all show the same spectrum (they only differ by their phase, i.e. their preferred direction), the spectrum of FS neurons are widely distributed, with many individual neurons showing high power in one of the Fourier components, resulting in symmetries in their tuning curves.

We reanalyzed the dataset of [23], composed of freely moving mouse recordings in the PoSub (n = 1529 pyramidal excitatory and n = 420 FS inhibitory cells) as well as in ADN (n = 102 ADN HD cells). As shown in [23], PoSub-FS cells have 1,2 or 3-folded tuning curves, as assessed by maximal Fourier power in their spectrum for the 1st, 2nd or 3rd component (Fig. 1A). The i-th Fourier power measures the variance explained by the i-th Fourier component, i.e how much a particular rotational frequency describes the tuning curve.

**Figure 1.**
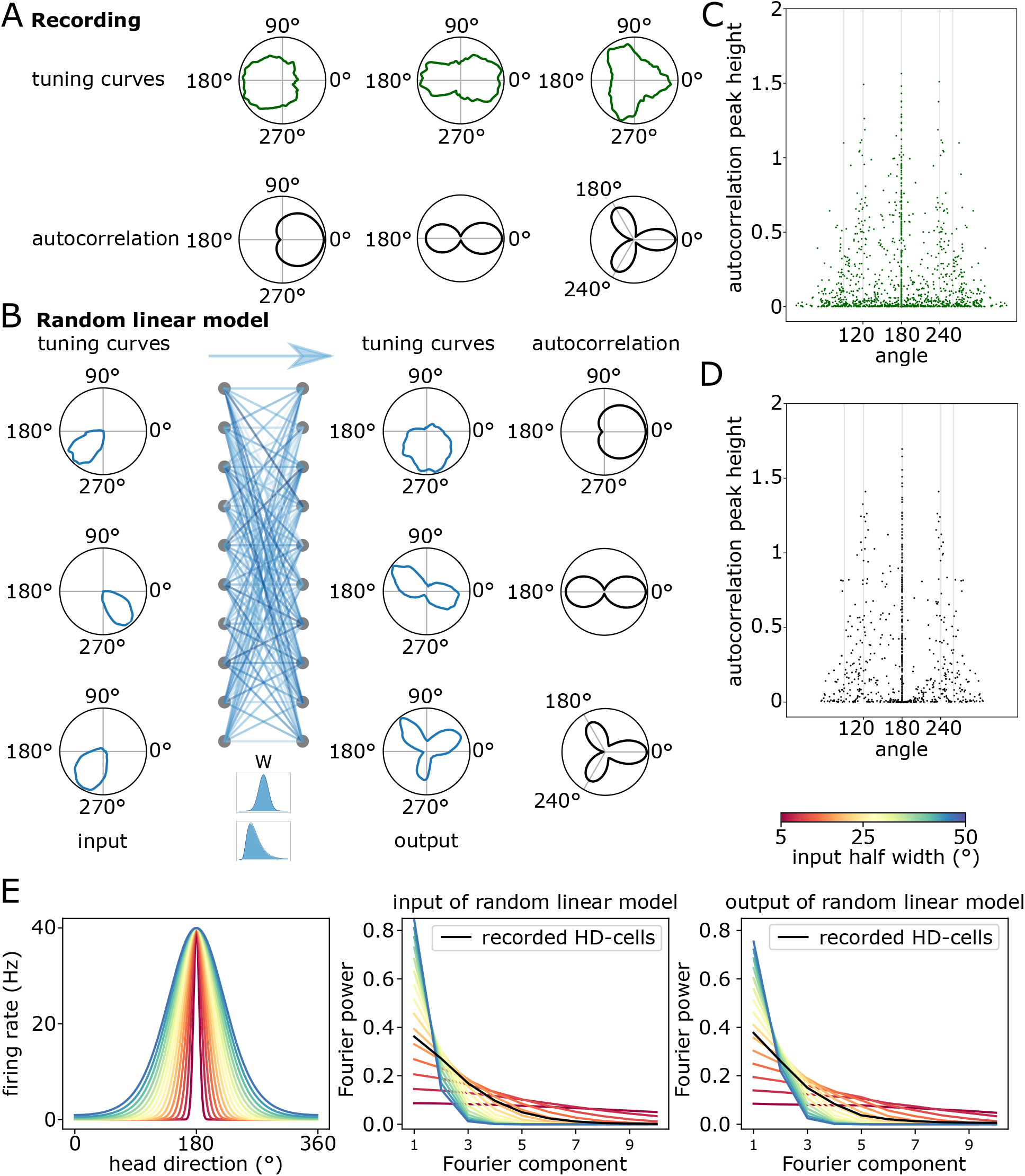
Symmetrical tuning curves in cortical fast-spiking neurons and how they emerge from random connectivity in a linear regime. A) Example FS cell tuning curves recorded from the head-direction cortex (top) and their autocorrelograms in feature space (bottom). B) Emergence of symmetries from random feed-forward connections. Left, example head-direction cell tuning curves recorded in the antero-dorsal nuclei of the thalamus. Middle, connectivity with either a normal or log-normal distribution. Right, tuning curves resulting from random linear combinations of input HD tuning curves, along with their auto-correlations C) Persistence diagram of tuning curves peaks, revealing symmetries at 180 and 120 degrees. D) the persistence diagram for 420 randomly generated tuning curves (As in C). Note the qualitative similarities with actual data shown in C. E) FS and excitatory cells share similar Fourier power in data and simulation. Left, canonical tuning curves of varying width. Middle, Fourier power spectrum of the canonical tuning curves, each tuning curves was normalized. Left, the mean Fourier power of the normalized random linear combinations of canonical input tuning curves. For each half-width we used 500 canonical tuning curve with uniformly distributed preferred distribution as input to the random linear model.

Cortical FS neurons receive inputs form local excitatory cells, but whether these inputs are organized is still a matter of debate. They can be considered as operating in a linear regime [24] and thus their activity may be captured by linear models of input integration. Interestingly, we observe similar symmetries in recorded FS cells and in a random linear model, combining canonical PoSub-HD tuning curves (Fig. 1 B). Indeed, the auto-correlation of the recorded (Fig. 1 C) and simulated (Fig. 1 D) tuning curves show dominant peaks at 120, 180, 240 degrees. While output FS cells tuning curves have more unspecific tuning curves, the mean Fourier power spectrum is shared between the input and output (Fig. 1 E). We next sought to establish a theoretical model accounting for the emergence of these symmetries in a random linear model.

### Distribution of tuning curves of linear neurons

#### General case

The response rate of a neuron relative to its inputs is a complex function of synaptic weights, membrane dynamics, and non-linear processes. Here, we sought to determine the average response of neurons to an encoded feature *θ* (i.e. its tuning) as a function of the tuning of its inputs and their synaptic weights. At first, we assume that the response of the neuron is linear and free of noise. In this case, the tuning curve is 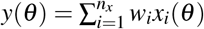, where *x* are the tuning curves of presynaptic neurons and *θ* is the feature.

What is the resulting distribution of tuning curve of output neurons? We first present the case of a linear combination with normally distributed weights *W* (mean *μ*_*W*_ and standard variation *σ*_*W*_). Tuning curves are discretized in *m*_*θ*_ points and this discretization is as fine as needed to preserve all the information of the tuning curves. The exact distribution of tuning curves can be derived by projecting the input tuning curves in their natural basis i.e by decomposing them with a singular value decomposition (SVD). The *n*_*x*_ *− by − m*_*θ*_ tuning curve matrix X is expressed as *X* = *USV*^*T*^. To determine the statistics of the output tuning curve *y* = *wX*, one can project *y* onto the singular vectors of *X*. Let us write *d* = *rank*(*X*), i.e d counts the number of non-zero eigenvalues in X. The projection onto the *j* = *d* + 1, …, *m*_*θ*_ vectors is 0. It is enough to know the value of the first d projection {*ŷ*_1_, …*ŷ*_*d*_} to fully reconstruct the tuning curve in the feature space: 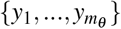. The distribution of tuning curves can be therefore equivalently described either in feature space or singular component space. The projection of *y* onto the jth singular vector is

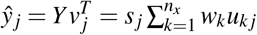

This is an affine transformation of the random multivariate normal variable *W* by *US*_(:,1:*d*)_. As *US*_(:,1:*d*)_ forms an orthogonal basis, the *ŷ*_*j*_ are independent (up to *j* = *d*). Specifically, the projection of a given tuning curve on the *j*th singular vector of its inputs only depends on the *j*th left singular component (or “score”) and the statistics of the synaptic weights. As *w*_*k*_ are independent samples from a normal probability distribution, the random variable *ŷ*_*j*_ is itself a normal distribution *𝒩* (*μ*_*j*_, *σ*_*j*_) where

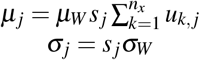

In summary, and quite intuitively, tuning curves of randomly connected linear neurons are constrained to lie in the subspace spanned by the singular component of their input tuning curves. They are distributed in this subspace according to a multivariate normal distribution with diagonal covariance. The weight standard deviation and input singular value *j* control the variance of *ŷ*_*j*_. This result is still valid for non-normally distributed weights in the limit of large number of input excitatory cells *n*_*x*_ by virtue of a generalized version of the central limit theorem (the Lindeberg-Feller theorem) (see methods and supplementary materials for proof). For example one might use only positive (excitatory) weights following a log-normal distribution. As shown below, these approximations are sufficient to account for the response of inhibitory cells in the cortex.

#### Tuning curves relative to a periodic variable

Periodic representations are ubiquitous in the brain, from orientation selectivity in the visual system to HD cells in the spatial navigation system. Remarkably, periodic tuning curves have right singular vectors that considerably simplify the understanding of their distribution. For a set of ideally distributed bell-shape curves (specifically, von Mises distributions with equally separated maxima), the set of singular vectors is a Fourier basis. Each odd component is a harmonic of the first vector, and each even component a 90 degree phase-shifted version of the preceding component, i.e. the natural orthogonal vector for a given harmonic (Fig. 2A). This is also approximately the case when the SVD is applied to a set of PoSub-HD cell tuning curves (Fig. 2B). The resulting singular vectors are qualitatively similar to the ideal case. We will therefore assume that tuning curve singular vectors can be identified with a Fourier basis.

**Figure 2.**
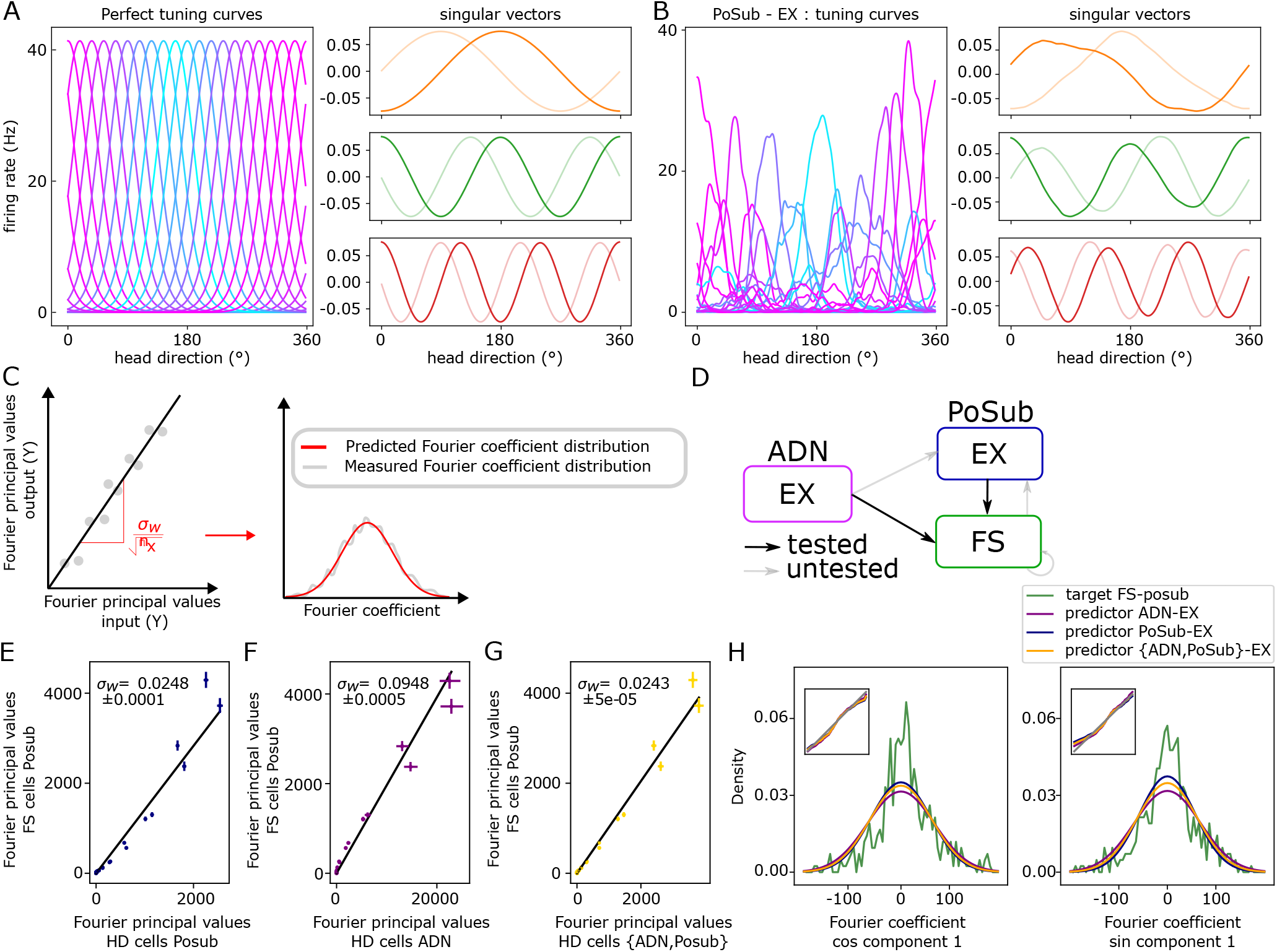
The distribution of FS cell Fourier coefficient, as revealed by SVD decomposition of tuning curves, is predicted by a random connectivity model. A) Left, example HD cell tuning curves. Right, first six singular vectors, grouped in pairs, of the tuning curves. B) Same as A for simulated HD cells, with homogeneously distributed preferred directions. In the ideal case, the set of singular vectors is equivalent to a Fourier basis. C) We predict that Fourier coefficient distribution are parametrized by the variance of synaptic weights *σ*_*w*_ (See text). We tested goodness-to-fit between actual and theoretical coefficient distribution. D) Anatomical diagram of the thalamocortical HD circuit. We tested predictions with inputs from either thalamus or local HD neurons to FS cells. E-G) Relationship between excitatory input and FS cell spectra in three cases: from PoSub (E), ADN (F), or both (G). H) Comparison of actual and predicted Fourier coefficient distribution for the first Fourier component.

Since the right singular vectors of inputs match with their Fourier basis, the distribution of the projection on the right singular vectors provides a multivariate distribution of output Fourier coefficients.

This multivariate normal distribution explains why symmetrical tuning curves emerge in the case of a linear model of input summation. From this distribution we can deduce the distribution of Fourier power 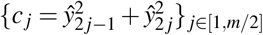, as the singular values of 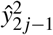 and 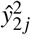 are approximately equal: *s* (*j*) := *s*_2*j*_ *≈ s*_2*j* − 1_. The distribution of a sum of squared independent normal variable with same variance is given by a scaled non-central chi-square of non-centrality parameter 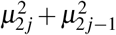 and degree of freedom 2:

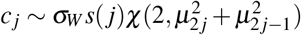

Importantly, unlike the precisely tuned input neurons which all show the same spectrum, each Fourier power of the tuning curves in the random linear model is an independent sample of a non-central chi-square distribution. Hence, although the power for the 1st component has a higher variance than the other components, it is possible that another component has a much larger power (Fig. S1), resulting in a tuning curve that shows strong symmetry at the corresponding frequency. Since the distribution of these non-central chi-square variables are controlled by the input power spectrum, the number of cells with a certain symmetry will be related to the power of each component in the input tuning curves (Fig. S1). In conclusion, the more energy in the input tuning curves (more selective) the higher the probability to linearly generate with random weights tuning curves with high-frequency symmetries.

In conclusion, symmetrical tuning can emerge from a random linear sum of narrowly tuned tuning curves. In particular, this model provides an exact formulation of the relationship between input tuning spectrum and the distribution of output tuning curves.

### Fit to experimental data

To assess the validity of the random linear model, it must be tested against the data. Specifically, this entails the comparison of the predicted multivariate distribution of tuning curve spectrum with the observed distribution. However, such statistical test is difficult. First a direct maximum-likelihood estimate of the weight distribution parameters (i.e. mean and standard deviation) is very sensitive to noise in singular vectors associated with low singular value (i.e. at high frequency, see supplementary materials). Second, as mentioned above, the Fourier basis is only an approximation of the true singular vector, hindering a rigorous test against the data. Here, we show however that representational-similarity matrices (RSM) of the input and output tuning curves must be proportional in the linear model. For the modelling of experimental noise and direct statistical tests see the supplementary materials.

#### Testing the model with representational-similarity matrices

Under the random linear model, the RSMs of the input and output tuning curves are proportional in the limit of a large number of recorded cells:

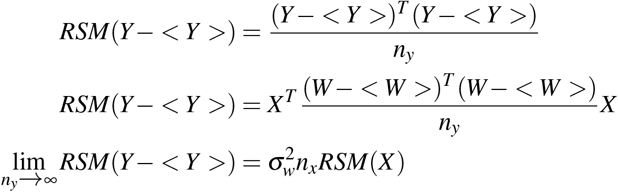

This result shows two key and testable predictions of the linear model. First, RSMs of the linear model input and output must be proportional, and the constant of proportionality varies with the variance of the weights. In other words, the slope of the fitting line provides the standard deviation of the weights, so that it is possible to compare the actual distribution of the tuning curves Fourier coefficients to the predicted normal distribution (parameterized by the measured variance) in the random linear model (Fig. 2C).

The convergence of the limit on *n*_*y*_ is quite slow and noisy estimate of the weight covariance matrix non-diagonal elements hinders model fitting. One solution is to project both RSMs on the shared right singular basis, here assumed to be the Fourier basis:

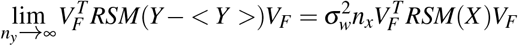

After projection, non-diagonal terms are zero in the limit so that the linear model can be tested against the diagonal elements only: 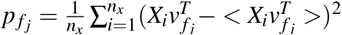. These diagonal elements are exactly the variance of the Fourier coefficients and will be referred hereafter to as the Fourier principal values.

We tested the random linear model for the following input-output ensembles in the experimental data: *PoSub → FS, ADN → FS*, and {*ADN, PoSub*} *→ FS* (Fig. 2D; RSMs are shown in Fig. S2). First we measured the goodness of fit of the RSM proportionality as assessed by the coefficient of determination: *R*^2^ = 0.97 *±* 0.006 (Fig. 2E), *R*2^=^0.957 *±* 0.021 (Fig. 2F) and *R*^2^ = 0.989 *±* 0.05 (Fig. 2G) for *PoSub → FS, ADN → FS*, and {*ADN, PoSub*} *→ FS*, respectively. Similar goodness of fit was obtained when using the full RSM matrix projected on the Fourier basis: *R*^2^ = 0.92 for ADN, *R*2 = 0.86 for PoSub.

We then compared the distribution of tuning curve Fourier coefficients to their normal distribution (Fig. 2 H). The fit was good for large number of components (first component presented in Fig. 2 H) but degraded at high frequency (Fig. S2). Error bars for the Fourier principal values were computed with a bootstrap procedure of Fourier principal values (Fig. S3 and methods).

#### Symmetries emerge for large weight variance

The expressiveness of the random linear model increases with the weight standard deviation (Fig. 3 A). In the case *σ*_*W*_ *<< μ*_*W*_, the proportions *{n*_*j*_*}* of folded cells are dictated by the homogeneity of the input tuning curves. Indeed in the limit *σ*_*W*_ *→* 0 each output tuning curve is the same: 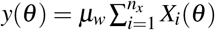. They simply reflect the sum of the input tuning curves. In the case where *σ*_*W*_ *>> μ*_*W*_, the proportion of folded cells increase and symmetrical tuning curves appear (Fig. 3 A).

**Figure 3.**
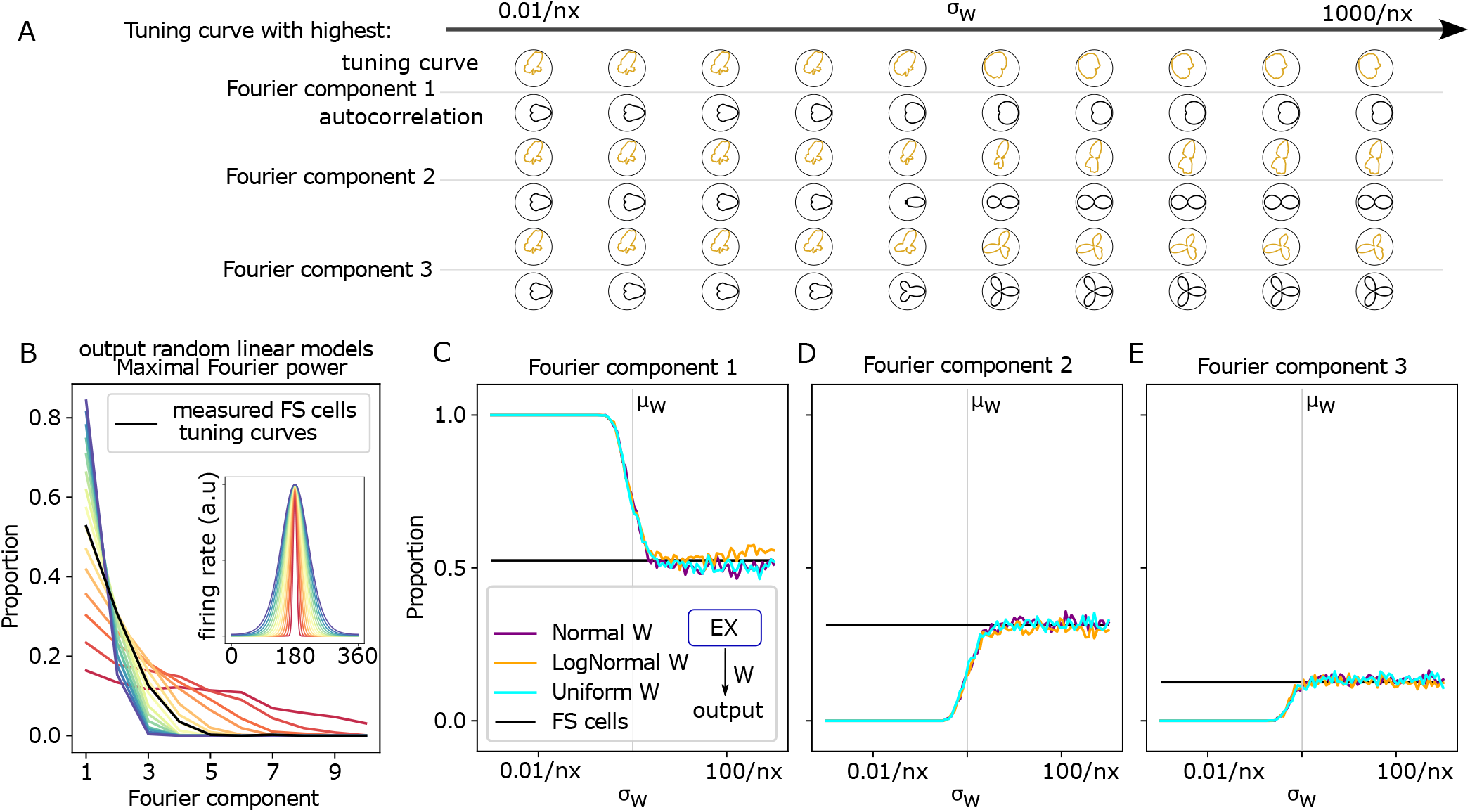
Symmetries in tuning curves emerge for large synaptic weight variance. A) Cell tuning curves with highest Fourier power in the first three components as a function of weight variance. B) Predicted ratios of tuning curves with highest power in the different Fourier components as a function of input tuning curve width (as in Fig. 1). Actual ratio of FS cells shown in black. C-E) Proportion of simulated cells with maximum power in the first (C) second (D) and third (E) Fourier component as a function of synaptic weight variance. Black lines display actual proportion of PoSub-FS cells.

More quantitatively, for random linear models characterized by *σ*_*W*_ *>> μ*_*W*_ and identical rotated tuning curves as inputs, the finer the input tuning curve half-width, the higher the number of output units showing high frequency symmetries (Fig. 3 B).

Next, we simulated the random linear model {*ADN, PoSub*} *→ FS* i.e with experimentally measured tuning curves as input. The proportion of n-folded output units converges to a limit value as the weight variance increases (Fig. 3 C,D,E). Remarkably this limit corresponds exactly to the measured proportion of PoSub-FS cells.

We provide an analytical derivation of these regimes and asymptotic values in the methods.

### Generalization to non-linear output units

We have so far focused on modeling the responses of PoSub-FS cells, which are well captured by a linear model. However, excitatory cells have non-linear responses. Can a simple model of non-linear output (Fig 4 A) reproduce the range of observed tuning curves across cell types in the HD system under the hypothesis of random connectivity? To address this question, we need to first determine the tuning properties that are uniquely associated with each cell class.

**Figure 4.**
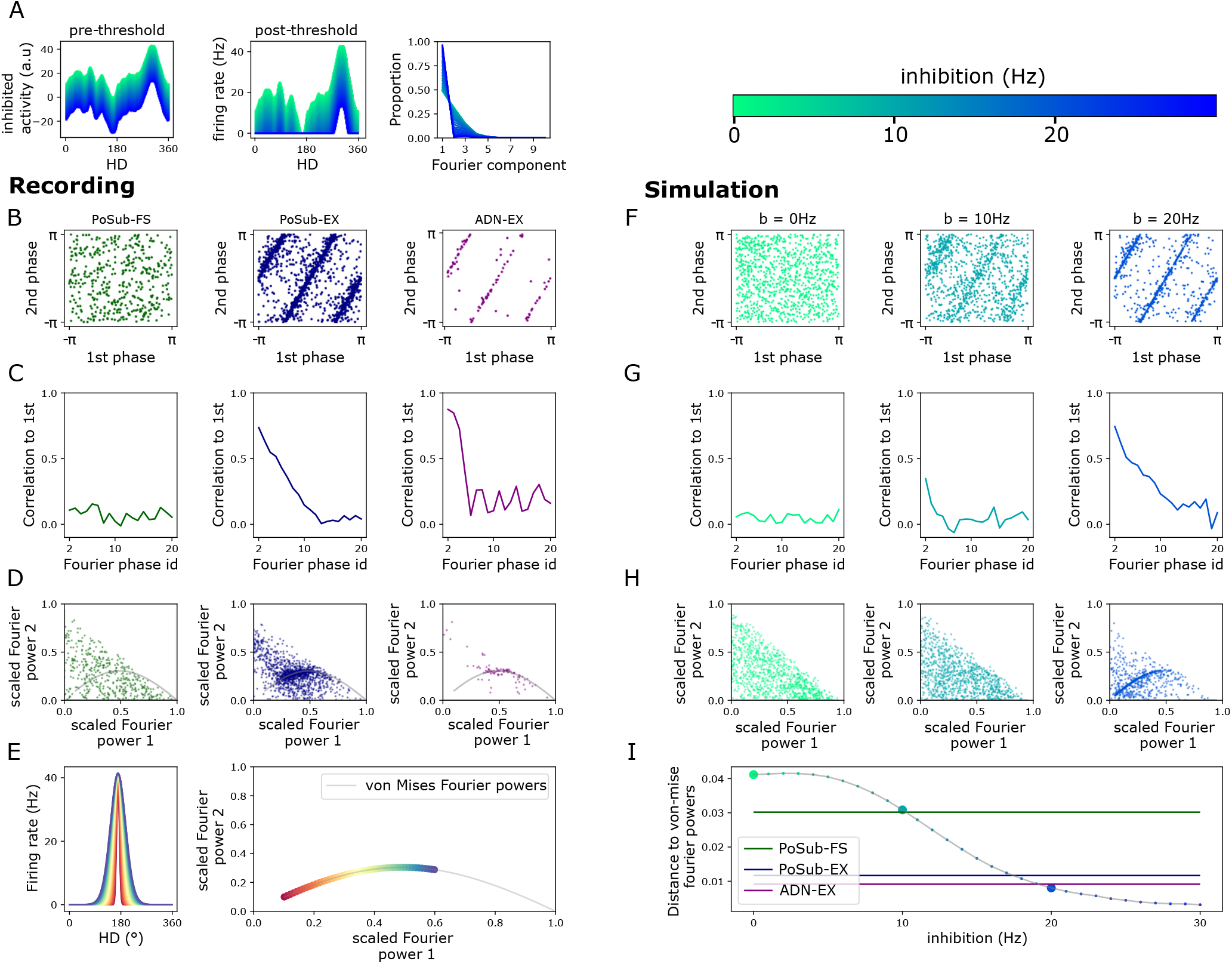
Random connectivity and homogeneous inhibition explains a continuum of output tuning curves. A) Iceberg effect. Left: subtractive inhibitions of a simulated FS tuning curve. Middle: tuning curves after relu composition. Right: proportion of simulated cells with maximum power in each Fourier components. B-E) Tuning curve properties of excitatory and inhibitory cells in the HD system. B) Phase-phase relationship of the first two components. C) Phase correlation between 1st component and higher component phases. D) Power-power relationship for the first two Fourier components. Line displays power-power relationship for ideal von Mises’ tuning curves (as explained in E). F-H) Tuning curve properties of simulated cells for various levels of additional inhibition, as in B-D). I) Mean distance of cells to the von-mise Fourier line plotted in E) as a function of the inhibition.

Excitatory and FS cells of the HD system differ in one key aspect: while the phases of Fourier components are independent in the case of FS cells, they are all interdependent for excitatory neurons. Specifically, the Fourier phase of the k-th component is related to the phase of the 1st component: *ϕ* (*k*) = *kϕ* (1) (Fig 4 B). This relationship is predicted by the Fourier decomposition of von Mises tuning curves (see methods). The same relationship in phases was observed in ADN- and PoSub-HD tuning curves. The phase-phase correlation disappeared at higher frequency (typically above the 10th component), as the tuning curves have necessarily limited resolution, at least because of the experimental limitations (i.e. accurate measurement of animal’s HD) (Fig 4 C).

Within the excitatory cell type of the thalamocortical HD system, the tuning curves of ADN- and PoSub-HD cells show one striking difference on the population level. While ADN-HD cells are not too different from the canonical response of ideal HD cells (i.e. bell shape and uni-modal tuning curve), many PoSub excitatory cells present multipeak tuning curves. This can be seen in the distribution of normalized power for the first two Fourier components (Fig 4 D). Ideal tuning curves, modeled as von Mises distribution of various angular widths, lie on a particular curve of the power-power plot (Fig 4 E). ADN cells are highly concentrated along this curve while PoSub excitatory neurons are more broadly distributed (Fig 4 D). Some PoSub neurons even had a second Fourier power larger than the first, corresponding to a two-fold symmetry of their tuning curves.

Remarkably, these two properties distinguishing ADN and PoSub neurons where all captured by a parsimonious non-linear random model [25]. Here, we assume that the tuning curve of a non-linear unit is: *Y* = *relu*(*WX − b*) where *W* captures both excitatory and inhibitory weight inputs, and *b* the addition of background inhibition (which does not necessarily correspond to physiological inhibition but could, for example, be related to higher spiking threshold). When no background inhibition is present and the unit integrates inputs from a sufficiently large number of input neurons, the total input to the unit is always suprathreshold, that is operates in the linear regime as in the previously studied case. When background inhibition exceeds the minimal of total input, the output response becomes non-linear. The progressive increase of this background inhibition leads to the shift from a multi-peak to single peak tuning curve. The transition between the linear and non-linear phase was associated with the progressive convergence of Fourier component phase (Fig 4 F,G) and power (Fig 4 H,I) to the distributions predicted by the von Mises model. The effect of background inhibition on response selectivity is a classic example of the “iceberg” effect [25, 26, 27] (Fig. 4 A).

### Random linear model predicts the effect of optogenetic manipulation of thalamic inputs to cortical neurons

In [23], the firing of ADN-HD cells was selectively disinhibited with optogenetics while recording in ADN or the PoSub. This gain manipulation did not change the tuning width in both ADN and PoSub, resulting in a purely multiplicative scaling of the excitatory HD cell tuning curves (Fig. 5 A).

**Figure 5.**
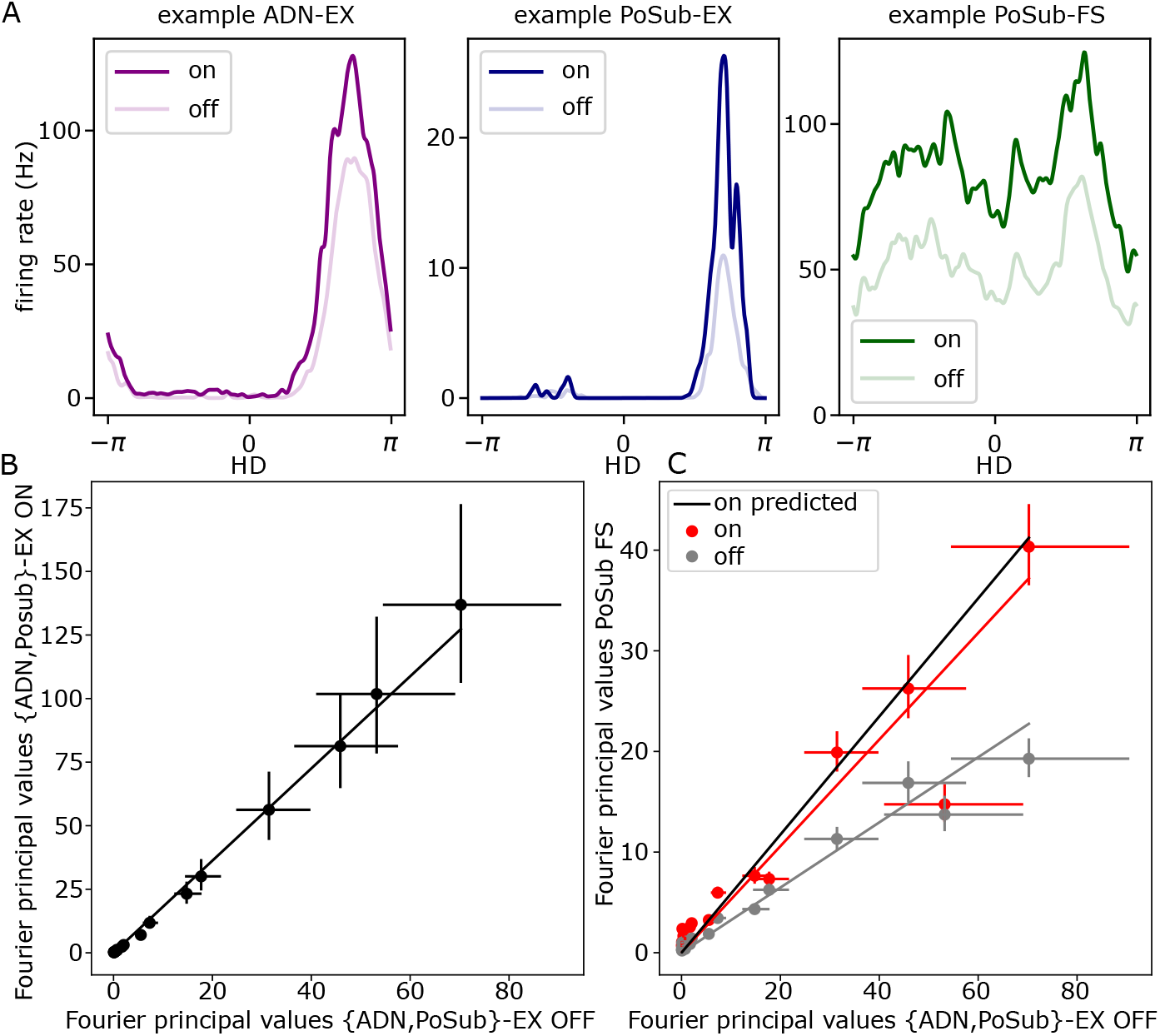
Predictions of the Random linear model are confirmed by an optogenetic manipulation. Tuning curves are multiplicatively increased during an optogenetic experiment (A), consequently all their Fourier principal values are scaled by a similar factor (B). For the random linear model this shared increase in variance in the input population should lead to a proportional increase in variance in the output, FS cells, population. Consequently the Fourier principal values of the FS cells population in the ON state should still be proportional to the excitatory cells Fourier principal values in the OFF state but with an increased multiplicative constant that matches the increase in the excitatory cells population (C).

The random linear model makes a clear prediction concerning this effect: a multiplicative gain of *α* for PoSub-HD and ADN-HD cell tuning curves is equivalent to a scaling of the input weights of FS cell standard deviations by the same factor *α*. Therefore we should have the following relationship between the Fourier principal values:

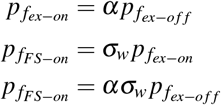

where *ex* and *FS* indexes refer to PoSub excitatory HD cells and FS cells, respectively.

We estimated the scaling factor *α* of the PoSub-ex HD Fourier principal values between ON and OFF conditions, and found *α* = 1.346 *±* 0.1 (Fig. 5 B). We then estimated the weight standard deviation *σ*_*w*_ = 0.569 *±* 0.034 based on the OFF excitatory and FS cell RSMs (Fig. 4 C). *alpha* and *σ*_*w*_ predicts the slope of the ON PoSub-FS cells Fourier principal values against the OFF PoSub-ex HD cells Fourier principal values. The estimated slope: *β* = 0.728 *±* 0.043 was within the error bar of the predicted one: 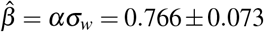 (Fig. 4 C). Error bars were computed by propagating the bootstrap estimates of measurement errors on the Fourier principal values (Fig. S3).

## Application to a different brain region

The random connectivity model is agnostic to the nature of the neuronal correlates. Here, we show that our analysis accounts for the tuning of FS cells in the hippocampus where a majority of excitatory cells are “place cells”, which fire for a specific position of the animal in the environment. To do so, we have re-analyzed a publicly available dataset [28] consisting of multichannel recording in CA1 while rats explored a linear track (1.6m or 2.0m in length). We computed place fields of every recorded cells in four rats, for a total of *n*_*x*_ = 348 putative pyramidal cells and *n*_*y*_ = 82 FS cells. The position was a 1d variable which we normalized between 0 and 1, independent of running directions. Both pyramidal and FS cells had tuning curves with single or multipeaks (Fig. 6A). As in the PoSub, both FS and pyramidal cells have symmetric tuning curves, with 2 or 3 peak as confirmed by the persistence scatter plots (Fig. 6B). In addition, some place cells tuning curve Fourier power spread along the von Mises line (Fig. 6 C) while FS cells did not (although more recorded cells would help to assess this effect).

**Figure 6.**
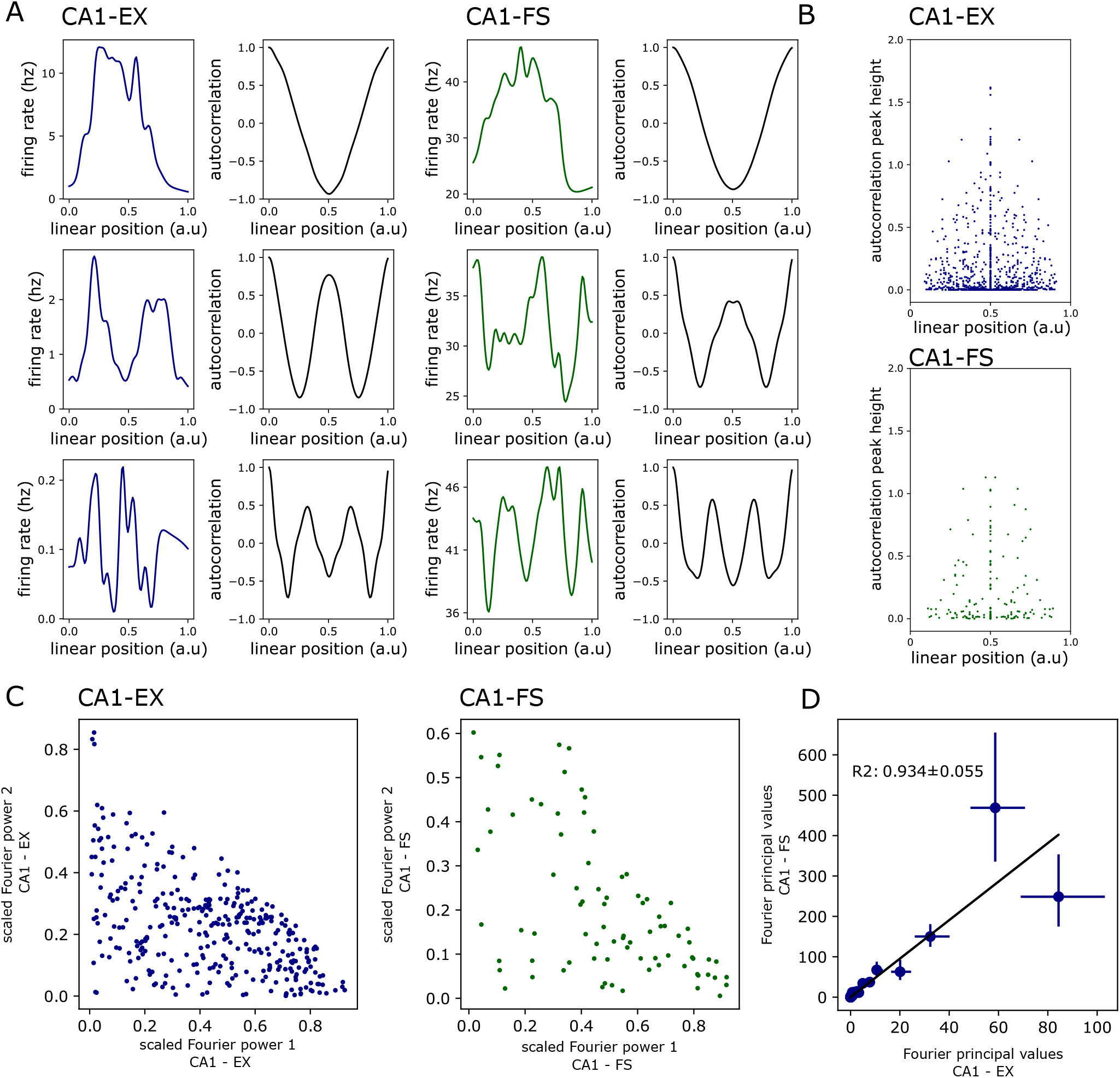
Random linear model for 1d linear tuning curves of cells in CA1. A) Left: tuning curve of CA1 excitatory cells (blue), with their autocorrelation (black) Right: tuning curve of CA1 FS cells (green) and their autocorrelation. B) Persistence diagram of the excitatory cells tuning curve (top) and FS cells tuning curve (bottom) C) Scatter plots of the first and second Fourier power divided by the sum of all Fourier power, for each tuning curves. Left: CA1 excitatory cell tuning curve, right: CA1 FS cell tuning curve D) We test if FS cells tuning curve can be generated from a random linear combination of excitatory cell tuning curves. Scatter plot of the Fourier principal values with the error bar estimated from a bootstrap strategy and propagation of uncertainty as for the HD dataset.

We tested if place fields of CA1-FS cells could be explained by a random projection from CA1 pyramidal cells. Fourier principal values of CA1-FS cells were well predicted by pyramidal cell Fourier principal values (*R*2 = 0.934 *±* 0.055, Fig. 6 D). Error bars and uncertainty propagation were estimated as previously. The first and second Fourier principal values (cos and sin of frequency one) were slightly off the prediction, either due to the limited number of cells per animal or highlighting a distinct spatial phase distribution in the pyramidal and FS population.

## Discussion

We found that randomly connected feed-forward networks can account for the diversity of tuning curves in the cortical stage of the head-direction system. In the linear case, random summation of input units showing ideal tuning curves leads to the emergence of symmetries in the output tuning curves. The distribution of symmetries correspond to the spectrum of the input tuning curves, as observed for PoSub-FS neurons relative to local excitatory cells [23]. We have derived an exact analytical solution, predicting the distribution of different symmetries in the FS cell population. The variance of the input weights is the key parameter of the model and symmetries appear when the variance exceeds a certain threshold. We then generalized the model to non-linear outputs. Varying background inhibition (or non-linearity threshold) is sufficient to reproduce in simulation all measured tuning curves in the Posub, including single and multipeaked PoSub excitatory tuning curves. We conclude that the distribution of PoSub HD tuning curves is compatible with a model of random connectivity for both PoSub excitatory inputs to local FS neurons and ADN inputs to PoSub neurons. Hence, these results support the view of an economical strategy for thalamocortical network wiring, which nevertheless supports reliable propagation of encoding properties.

The model predicted well the effect of artificial input disinhibition on FS tuning curves. Furthermore, it accounted also for the tuning curves of FS cells in the hippocampus in relation this time to place coding. Together these results show that the model can be generalized to a broad class of brain system.

The present study could not provide any direct test of underlying connectivity. Previous work which sought to address the relationship between connectivity and function reach similar conclusions for cortical FS cells, specifically that, in the visual cortex, excitatory inputs onto FS cells was largely determined by anatomical proximity rather than tuning properties [15, 17]. Furthermore, the random model is sufficient, but not necessary, to account for the emergence of observed tuning curves. Whether more complex organization principles govern cortical neurons at the population level is still an open question.

The present model does not take into account recurrent connections within the PoSub. While connections between excitatory neurons is rare in the PoSub, especially in the superficial layers that receive inputs from ADN, excitatory and FS cells are mutually connected [29]. We assumed in the non-linear model that background inhibition was untuned to the features. It is likely that the tuning of FS cells, as observed and modelled here, play an additional role in shaping excitatory cell tuning curves. In addition, FS cells show high probability of mutual connections and it is possible that this reciprocal inhibition further influence the tuning of FS cells. Accounting for these recurrent connections within the PoSub and their potential consequences for neuronal coding will be the focus of future work.

This model relies on the knowledge of cell’s tuning curves. This applies well to primary sensory systems in which tuning is generally well understood but may prove difficult in other areas where cell tuning to task variables is complex [30], if not unknown. However, it could be generalized to all systems in which population activity is of low dimensionality and intrinsic feature can be determined from the population activity and not necessarily from external attributes [31, 32, 33, 34, 35]. Additional work is necessary to address the tuning of FS cells in a network where excitatory neurons encode multiple variables [30].

### Emergence of symmetries

Our work defined a symmetrical tuning curve by the maximal power of its tuning curve Fourier spectrum. This definition would make any tuning curve necessarily symmetric for some frequency, up to the sampling resolution of the feature. Our analytical work shows that, in the linear model, only symmetries at small frequency should emerge, as observed in the data [23]. FS cells have often been considered broadly tuned to features [13, 14, 15, 16, 17, 18, 19, 20, 21, 22], with some being narrowly tuned [36]. Here, we thus show that this broad range of tuning can emerge from a random summation of tuned local excitatory cells. A key take-away from our analytical framework is that Fourier coefficients are independent. Hence, resulting output units show symmetries at frequencies that are intrinsic to the shape of the input tuning curves (with narrower tuning curves conveying higher frequencies) although input tuning curves do not necessarily show symmetries.

These symmetries emerge altogether above a certain threshold of input weight variance. This threshold is in the order of, but not exactly equal to the weigh distribution mean. Below this threshold, output units reflect input tuning curves and are all dominated by the 1st Fourier component. It is important to note that connection weights may not be the only source of variance. The model described here is equivalent to a system where all weights have equal strength but input unit show various level of peak firing rate (or excitability). In fact, both properties, connection weights and cell’s excitability, may be broadly distributed [37].

In the non-linear model, high level of background inhibition makes the output strongly non-linear and results in uni-modal tuning curves that can be even narrower than input tuning curves. This is usually referred to as the “iceberg” effect [26, 27]. This would simply explain how PoSub-HD neurons become more narrowly tuned than their thalamic inputs. Interestingly, for intermediate level of inhibition, non-linear output units can be both selective and symmetrical, that is their tuning curves is multipeaked and the angular offset between these peaks is not random. This reflects the experimental observation where PoSub-EX cells are more multipeaked than ADN-EX neurons. Such symmetries in the HD tuning of excitatory cells have been observed in other areas of the brain’s navigation system [38, 39]. Whether these symmetries emerge from the nature of their inputs or are driven by external factors (e.g. vision) remains to be determined.

### Random linear model and learning

Contrary to the random model, the similarity between FS cells tuning curve and the Fourier basis may suggest that the network connectivity is optimized to extract this particular basis. Indeed, in certain class of models, the natural basis of input signals spontaneously emerges either through principal components analysis [40, 41, 42], independent components analysis [43] or maximization of mutual information [44, 45, 46]. However, output unit of the random model, which share striking similarities with FS cells, do not form an actual basis but rather a random expansion of this basis. Output units are characterized by random combination of different Fourier components with independent but non-identical normal distribution. Remarkably, the variance of each normal Fourier component distribution reflect the relative power of the input tuning curve Fourier spectrum. This would not be the case if the inputs were whitened, which would result in output unit tuned to any possible frequencies.

Random models are not a mere computational abstraction but, instead, may prove crucial for representation learning. Specifically, the variance of random weights when initializing artificial neural networks controls their post-training generalization [47] and training dynamics [48]. In the “lazy” learning regime, i.e high variance of the initial weight distribution, the information propagates well through the network, as observed in our data and model. This could potentially help learning to occur rapidly through all layers. Similarly, in recurrent neural networks, task learning may require minimal change from random connectivity, often spanning a low number of ranks of the connectivity matrix [12, 49, 50, 51]. Finally, in convolutional neural networks, randomly initialized filters are known to be maximally responding to 2D sinusoidal pattern at specific frequency, i.e they are sensitive to Gabors filter even without weight learning [52].

## Conclusion

Our work shows that the diversity of cortical tuning in primary sensory areas may result from random summation of inputs, either from local population or upstream neurons in the thalamus. This suggests an economical rather than precisely organized wiring principles for a large class of brain networks. These findings have important implications for our understanding of how brain areas connect to each other and how high-level representation emerge in thalamocortical networks.

## Acknowledgements

This project has received funding from the European Union’s Horizon 2020 research and innovation program under the Marie Skłodowska-Curie grant agreement No 945304. (P.O) ; a Canadian Research Chair in Systems Neuroscience, CIHR Project Grant 155957, NSERC Discovery Grant RGPIN-2018-04600, and the Canada-Israel Health Research Initiative, jointly funded by the Canadian Institutes of Health Research, the Israel Science Foundation, the International Development Research Centre, Canada and the Azrieli Foundation 108877-001 (A.P.); EMBO Long-Term Postdoctoral Fellowship and Sir Henry Wellcome Fellowship (A.J.D.).

## Author contributions statement

P.O. performed the analysis and modeling. A.D. recorded the data and made the observation of symmetrical FS-tuning curves. P.O. and A.P. wrote the manuscript. A.P. supervised the project.

## Methods

### Data

The dataset of recordings in the HD thalamocortical system is described in [23]. Briefly, in this dataset, the animal’s head-direction was monitored with a set of six cameras tracking markers glued on the animal’s hat (Optitrack, OR, USA) while the animal was foraging for food in a square arena. The system captured the 3-dimensional coordinates of all tracking markers, relative to the global coordinates, constant across the whole study, from which the head-direction was computed. In order to minimize the effect of animal’s velocity on HD tuning curves, the dataset was limited to the epochs when animal’s speed exceeded 2 cm/s for all analyses except the optogenetic experiment, where epoch duration (240 s in total) was too short to warrant further refinement. HD tuning curves were then computed as the ratio between histograms of spike count and total time spent in each direction in bins of 1 degree and smoothed with a Gaussian kernel of 3° s.d. The present results are independent of tuning curve smoothing: they could be reproduced using raw tuning curves (not shown).

The CA1 dataset is the HC-11 dataset [28] available on the CRCNS website ^1^. We focused our analysis on the linear track sessions.

### Analysis

All analyses were performed in Python 3.7 using standard libraries.

#### Autocorrelation

The autocorrelogram of a tuning curve *y* ∈ **R**^(^*m*_*θ*_), of mean *μ* and std *σ* is computed as:

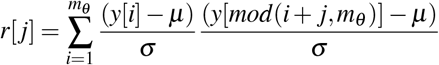

We then extract the peak in the autocorrelogram and compute the persistence of each peak, using function from the scipy package (scipy.signal.find_peaks and scipy.signal.peak_prominences, respectively).

#### Fourier analysis

In Fig. 2 and 5, tuning curves are projected on a discrete Fourier basis of size 2*n*_*f*_ = 2 × 40 evaluated over *m*_*θ*_ = 360 angles: *θ* ∈ {0,.., 359 × *π*/180}:

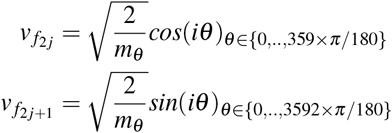

The factor 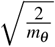 normalizes each vector of the basis. For Fig. 1E, the Fourier power spectrum of each tuning curve is normalized by their sum (equivalent to dividing the tuning curve by its variance).

The Fourier principal values of *X* (a set of tuning curves) are the variance of the set of Fourier coefficients:

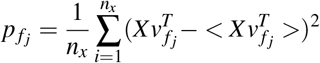

#### Normal distribution of projection on input right singular vector

Irrespective of the weight distribution, the central limit theorem states that the random variable 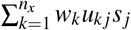 should converge to a normal distribution. However, the central limit theorem does not exactly applies here because of the non-randomness of the *u*_*k j*_*s* _*j*_. Instead, we prove in the supplementary materials that a generalized version of the central limit theorem holds in this case, by demonstrating that the sequence of random variable *w*_*k*_*u*_*k j*_*s* _*j*_ verifies a Lindberg condition which is sufficient for a central limit theorem to apply. This is due to the increase of the SVD spectrum {*s*_*j*_} as a function of the number of input neurons *n*_*x*_.

#### Effect of the mean and variance on the proportion of n-folded tuning curves

Let us rewrite each weight as *w* = *μ*_*w*_ + *νσ*_*w*_. Each tuning curve is then

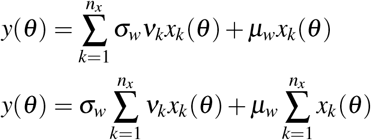

Tuning curves are thus expressed as the sum of two terms: 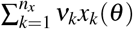 and 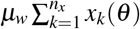. for normally distributed weights, the first term is independent of the weight mean and standard deviation; Therefore, it must remain constant constant when the standard variation increases (Fig. 3), with variation only depending on the random sample.

Let us note the Fourier coefficients of this term as *b*_*l*_: 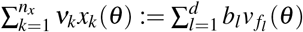.

Let us also note the Fourier coefficients of the sum of input tuning curves as *a*_*l*_: 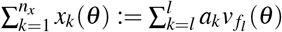

Consequently, the Fourier coefficients of the output tuning curves are: *ŷ*_*l*_ = *σ*_*w*_*b*_*l*_ + *μ*_*w*_*a*_*l*_

The Fourier power of the output tuning curves are: 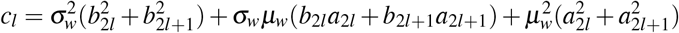

In two extreme limit of the ratio 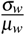, the Fourier power converge to a limit distribution:

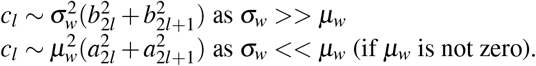

The proportion of cells with a certain maximal Fourier power is a statistic of the joint ratio of each Fourier power over the sum of all Fourier powers. The random variable associated with this ratio is:

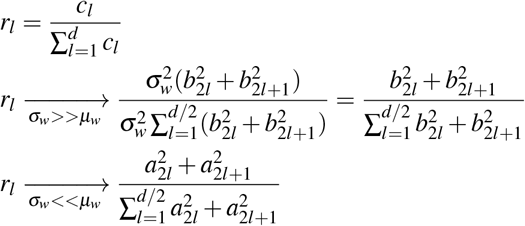

These terms become independent of the exact value of the mean and standard deviation which explains the asymptotic proportion in simulations for the normal random variable (Fig. 3 C). Similar derivations can be made for non-normally distributed weights as Fourier coefficients are normally distributed irrespective of the exact weight distribution, in the limit of a large number of input cells.

#### Modeling the effect of sampling on Fourier principal values

The sampling of neurons is necessarily limited by experimental constraints. How does it affect the statistics of the tuning? We propose to estimate the measurement error of tuning curves across all cell populations on the Fourier principal values *p* _*f j*_ by a bootstrap procedure. This uncertainty will allow us to get confidence intervals on the linear relationship fitted in the main paper (Figs. 2, 5), i.e the linear relationship between Fourier principal values of input and output tuning curves.

We randomly sub-sampled *n*_*s*_ tuning curves from the set of recorded tuning curves in each population and estimated the standard deviation of their Fourier power. By repeating this procedure a thousand times we obtained bootstrapped estimates of the distribution of these Fourier principal values. When sub-sampled with replacement these distributions are well fitted by Log-Normal distributions with similar log-standard deviation *σ*_*l*_ across Fourier component.

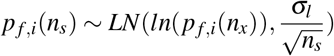

Where *p*_*f,i*_(*n*_*x*_) is the i-th Fourier principal value when using all *n*_*x*_ cells in the population. If no replacement was made, this noise estimate was still good for low *n*_*s*_ but degraded, as expected, when *n*_*s*_ approached the number of recorded cells, with the standard deviation falling faster than 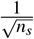. Example of fit for *n*_*s*_ = 100 are presented in Fig. S2. The scaling factor *σ*_*l*_ was found to be very similar across the three population (ADN-EX, Posub-EX, Posub-FS). For better precision we kept the individually estimated *σ*_*l*_ for each Fourier principal value.

The measurement error we make on each Fourier principal value will propagate to statistics obtained from them, notably the weight standard deviation and the R2-score. Hopefully we just observed that the measurement error in the experimentally measurable quantities is log-linear and therefore multiplicative. We can therefore propagate this error in the following way.. Let us define *ε*_*in*_(*n*_*s*_) and *ε*_*out*_(*n*_*s*_) the input and output principal values error term, respectively. We have that *ε*_*in*_(*n*_*s*_) *∼ LN*(0, *σ*_*l*_*/n*_*s*_), *ε*_*out*_(*n*_*s*_) *∼ LN*(0, *σ*_*l*_*/n*_*s*_).

The random linear model makes the predictions:

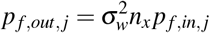

We rewrite each of the measured factor as a multiplication of the observed value and its multiplicative error term and apply a logarithm to both side of the equation:

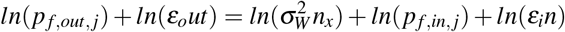

The terms *ln*(*ε*_*o*_*ut*) and *ln*(*ε*_*i*_*n*) are both normally distributed as 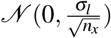 as explained above. The weight standard deviation is estimated by a linear regression as

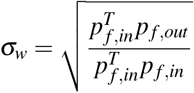

To propagate errors we rewrite the input Fourier principal values as 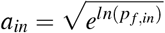, since we have a *±* error estimate on *ln*(*p*_*f,in*_), and similarly for the output Fourier principal values. *σ*_*w*_ is:

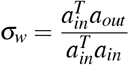

This formulation makes it possible to propagats the error from *ln*(*p*_*f,in*_) and *ln*(*p*_*f,out*_) to *σ*_*w*_. The measurements error computed with the Uncertainties package.^2^.

A similar sampling effect was found for the CA1 dataset [28]. Results can be found in Fig. S3.

#### von Mises model

In Fig. S6, we give a more complete overview of the Fourier phases of the experimentally recorded tuning curves. It is clear from the data that the phases of the FS cells are independent while the phases of excitatory cells are correlated. This dependence for excitatory cells can be understood through a Fourier decomposition of typical von Mises tuning curves:

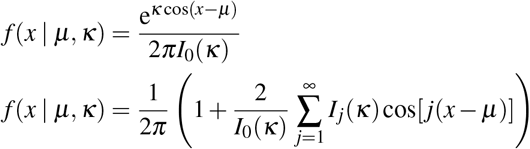

Where *μ, κ* parametrize von Mises tuning curves and *I*_*j*_(*κ*) is the j-th modified Bessel function of first order.

We used in simulation a slightly different but equivalent parameterization of the von Mises distribution so that the tuning curves had similar maximal firing rate.

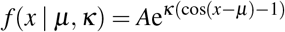

The Fourier decomposition of the von Mises tuning curve indicates that the *j − th* Fourier phase of a cell with preferred direction *ϕ* (1) (i.e. the phase of the first Fourier component) should be *ϕ* (*j*) = *jμ* = *jϕ* (1).

It also provides the set of Fourier power for perfect von Mises tuning curves: 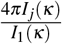, which defines the von Mises Fourier power line presented in the main figures (as a function of *κ*). We have highlighted that the 1st and 2nd Fourier components of excitatory cells seem to converge toward the 1st and 2nd components of von Mises tuning curve, in the data and in our simulation when we use a relu-nonlinearity.

Note that the Bessel functions are the set of solutions of:

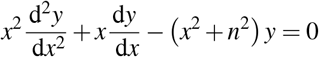

The existence of an underlying link between this property and the iceberg effect is left for future works.

## Footnotes

1 http://crcns.org/data-sets/hc/hc-11/about-hc-11.

2 Uncertainties: a Python package for calculations with uncertainties, Eric O. LEBIGOT

## Notes

### Competing Interest Statement

The authors have declared no competing interest.

