## Supplementary materials for "Signature of random connectivity in the distribution of neuronal tuning curves"

### Supplementary texts

#### Proof of the central limit theorem

The Lindberg-Feller central limit theorem is the following:
Suppose $X_1, ...X_n$ are independent random variable with mean 0 and finite variance $\sigma_j^2$. Let $s_n^2 = \sum_{k=1}^n \sigma_j^2$ denote the variance of their sum. If for every $\varepsilon > 0$,

$$\frac{1}{s_n^2}\sum_{j=1}^{n} E(X_j^2 1_{|X_j|>\varepsilon s_n}) \xrightarrow[n\to\infty]{} 0$$

then $S_n/s_n \to \mathcal{N}(0,1)$.

The theroem condition is called the Lindberg condition. Let us demonstrate the the limit holds, we define $X_j = W_j u_{jk} s_k - \mu_w u_{jk} s_k$ which are of mean 0 and finite variance. $s_n^2 = \sum_{k=1}^{n_x} \sigma_w^2 u_{jk}^2 s_k^2 = s_k^2 \sigma_w$

Note that $s_k^2$ is a function of $n_x$ monotonically growing to infinity.

Be $\varepsilon > 0$ The Lindberg condition left hand term can be written as:

$$\frac{1}{s_n^2}\sum_{j=1}^{n} E(X_j^2 1_{|X_j|>\varepsilon s_n}) = \frac{1}{\sigma_w}\sum_{j=1}^{n} u_{jk}^2 E((W_j-\mu_W)^2 1_{|W_j-\mu_W|>\varepsilon\sigma_w \frac{|s_k|}{|u_{jk}|}})$$

We have: $P(|W_j - \mu_W| > \varepsilon\sigma_w \frac{|s_k|}{|u_{jk}|}) \leq P(|W_j - \mu_W| > \varepsilon\sigma_w \frac{|s_k|}{max_j(|u_{jk}|)})$ which is independent of j.

Let us write $A_k = |W_j - \mu_W| > \varepsilon\sigma_w \frac{|s_k|}{max_j(|u_{jk}|)}$.

The weights have i.i.d distribution, we can therefore bound the left hand term by: $\frac{E((W-\mu_W)^2 1_{A_k} \sum_{j=1}^n u_{jk}^2)}{\sigma_w}$

Let us define $Z_n = (W-\mu_W)^2 1_{A_k} \sum_{j=1}^n u_{jk}^2)$ We have: $\sum_{j=1}^n u_{jk}^2 = 1$

Moreover $\frac{s_k}{max_j u_{jk}} \to \infty$ with $n_x$, indeed $max_j u_{jk} < 1$ as $\sum_j u_{jk}^2 = 1$ and $s_k \to \infty$ with $n_x$

Therefore we have $P(|W_j - \mu_W| > \varepsilon\sigma_w \frac{s_k}{u_{jk}}) \to 0$, and by definition of convergence by probability: $Z_{n_x} \xrightarrow{P} 0$.

Moreover, $Z_n \leq (W-\mu_w)^2$ and $E(((W-\mu_w))^2) \leq \infty$ so by dominated convergence theorem : $E(Z_n) \xrightarrow[n\to\infty]{} 0$.

Consequently we have $\frac{1}{s_n^2}\sum_{j=1}^n E(X_j^2 1_{|X_j|>\varepsilon s_n}) \to 0$. This was independent of $\varepsilon$ and consequently we have proven that the Lindberg condition holds which concludes the proof.

While this proves the normality of each singular value projection of the output tuning curves, we have still to show their independence. The independence comes from the fact that the covariance matrix is diagonal. A multivariate normal distribution with diagonal covariance has independent components. Therefore the singular value projection of the output tuning curves are indeed independent.

#### Maximum Likelihood estimates

Given a set of measured input $X$ and output $Y$ tuning curves and assuming $Y = WX$, the weights $W$ parameters $\sigma_W, \mu_W$ can be estimated by a maximum likelihood approach. These parameters should maximize the likelihood $L(Y,X)$ of having observed $(X,Y)$.

$$\hat{\mu}_W, \hat{\sigma}_W = argmax_{\mu_W,\sigma_W}(\prod_{i=1}^n pdf(Y_i))$$

$$\hat{\mu}_W = \frac{1}{\sum_{j=1}^{m} a_j^2}\left(\sum_{j=1}^{m} a_j \frac{\frac{1}{n_y}\sum_{i=1}^{n_y}\hat{y}_{i,j}}{s_j}\right)$$
$$\hat{\sigma}_W = \sqrt{\frac{1}{mn_y}\sum_{j=1}^{m}\sum_{i=1}^{n_y}\left(\frac{\hat{y}_{i,j}-\hat{\mu}_W s_j a_j}{s_j}\right)^2}$$

Where $a_j = \sum_{k=1}^{n_x} u_{k,j}$.

A first drawbacks of these MLE estimate is that they will be very sensitive to noise in the Fourier coefficients of high frequency. Indeed if we model a little deviation $\varepsilon_j$ for each Fourier components: $\bar{y_{i,j}} = \hat{y}_{i,j} + \varepsilon_j$, the sensitivity of our estimates to these deviations are:

$$|\frac{\partial \hat{\mu}_W}{\partial \varepsilon_j}| = |\frac{a_j}{s_j \sum_{l=1}^{m} a_l^2}|$$
$$|\frac{\partial \hat{\sigma}_W}{\partial \varepsilon_j}| = |2\hat{\sigma}_W \frac{\sum_{i=1}^{n_y}(\hat{y}_{i,j}+\varepsilon_j-\hat{\mu}_W s_j a_j)(1+\frac{\partial \hat{\mu}_W}{\partial \varepsilon_j} s_j a_j)}{mn_y s_j^2}|$$

For Fourier components of high frequency, i.e where $s_j$ is small, these sensitivity will have very large values.

Second they assume that the SVD right singular vector of the input tuning curves are exactly known, which is not the case, simply because we don't record from all cells.

#### Likelihood ratio tests

In contrast to FS Fourier coefficients, PoSub-EX tuning curves Fourier coefficients should not be independently distributed from each other. Nonetheless the non-independence of the PoSub-EX tuning curve Fourier coefficients and independence of FS cell Fourier coefficients could not be statistically assessed when testing directly for the multivariate independence of the Fourier components. Such test of independence can be computed as a likelihood-ratio test with the null-hypothesis given by a null correlation matrix. The test statistic is then given by: $\Lambda = \frac{|\Sigma|}{\prod_{i=1}^{m}\Sigma_{ii}}$. In our hand this statistic was too weak to refute the multivariate independence of the PoSub-EX cells. To show this we computed the statistics at equal sampling size with the FS cells by subsampling the number of excitatory cells in the PoSub population. We also simulated multiple random linear projection from von Mises tuning curve with a random but uniform sets of prefered angles. This simulation was repeated for a varying amount of simulated excitatory inputs, and we report the test statistics in (suppFig. 5). FS test statistic was inside the distribution of PoSub-EX test statistics. Perfect random linear model would generate similar statistics for a wide range of number of input cells. This shows that this multivariate test for independence is in practice too weak to assert if the Fourier components are independent to each other. For this analysis we projected both the simulated and the experimentally measured set of cells on the Fourier basis, and the results was robust to the number of Fourier basis used.

### Non-independence of powers

The random linear models predicts independent non-central chi-square distribution for the Fourier power. But when we tested this independence with a correlation test it appeared not to be the case for FS cells $r = 0.64$, $p < 0.01$ (SuppFig 6). When using Normally distributed weights such correlation was absent in our simulations (as predicted by the theory), but once we departed from normality, such correlation could be generated, notably using Log-Normally distributed weights. We observed that such correlation is indeed driven by high-firing variance cells. Since the sum of Fourier power is proportional to the cell variance, cells with large variance necessarily will have large Fourier powers. In the main paper we therefore scatter plot the variance-free Fourier power in Fig. 4. The independence of FS cell Fourier powers irrespective of variance is observable in this plot.

#### Effect on cortical maps

We simulate the effect of the random linear model by summing excitatory cells spread randomly on a squared cortical sheet. The symmetry of local sub-population tuning curves, generated by a sum of excitatory cells tuning curves, are depicted in figure 8. As expected when the iceberg effect is applied on this sheet, the number of symmetries when can see emerging in the sub-populations increase with the inhibition strength. Note that all excitatory cells preserve their single-peak tuning curves, but have increase selectivity as the iceberg effect acts and the inhibition increases.

### Supplementary Figures

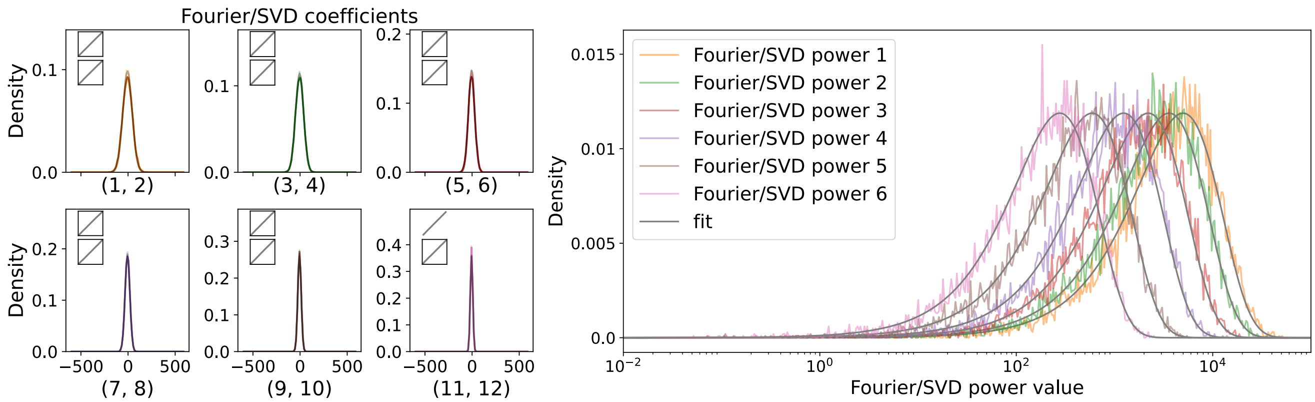


**Supplementary Figure 1.** Comparison of theory and simulation for a finite size random linear projection. Left: the distribution of each Fourier coefficients, right: the distribution of Fourier power. Predicted fits are displayed in black.

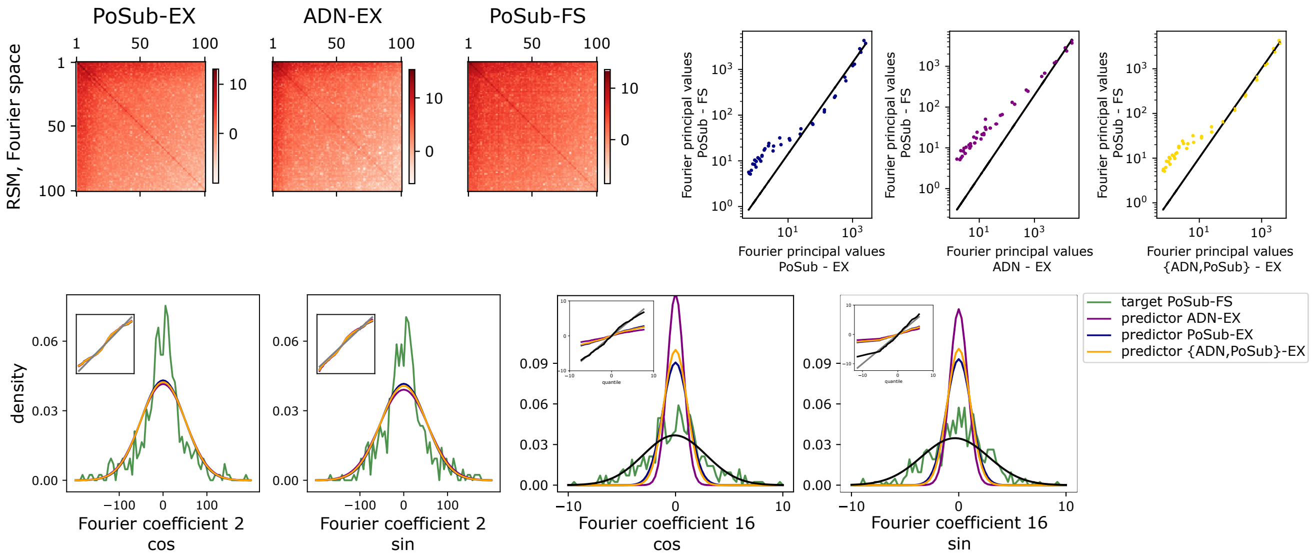


**Supplementary Figure 2.** Additional components for the fit of the random linear model of the HD dataset. Top left: RSM matrix for each sets of tuning curve. Each RSM is projected on the right and left with the Fourier basis and displayed in log scale. Top right: Fourier principal values with a logarithmic scale (40 Fourier components, the following Fourier components follow the same parallel trend to the fit line). Bottom: Fourier components 2 (well fitted) and 16 (poorly fitted).

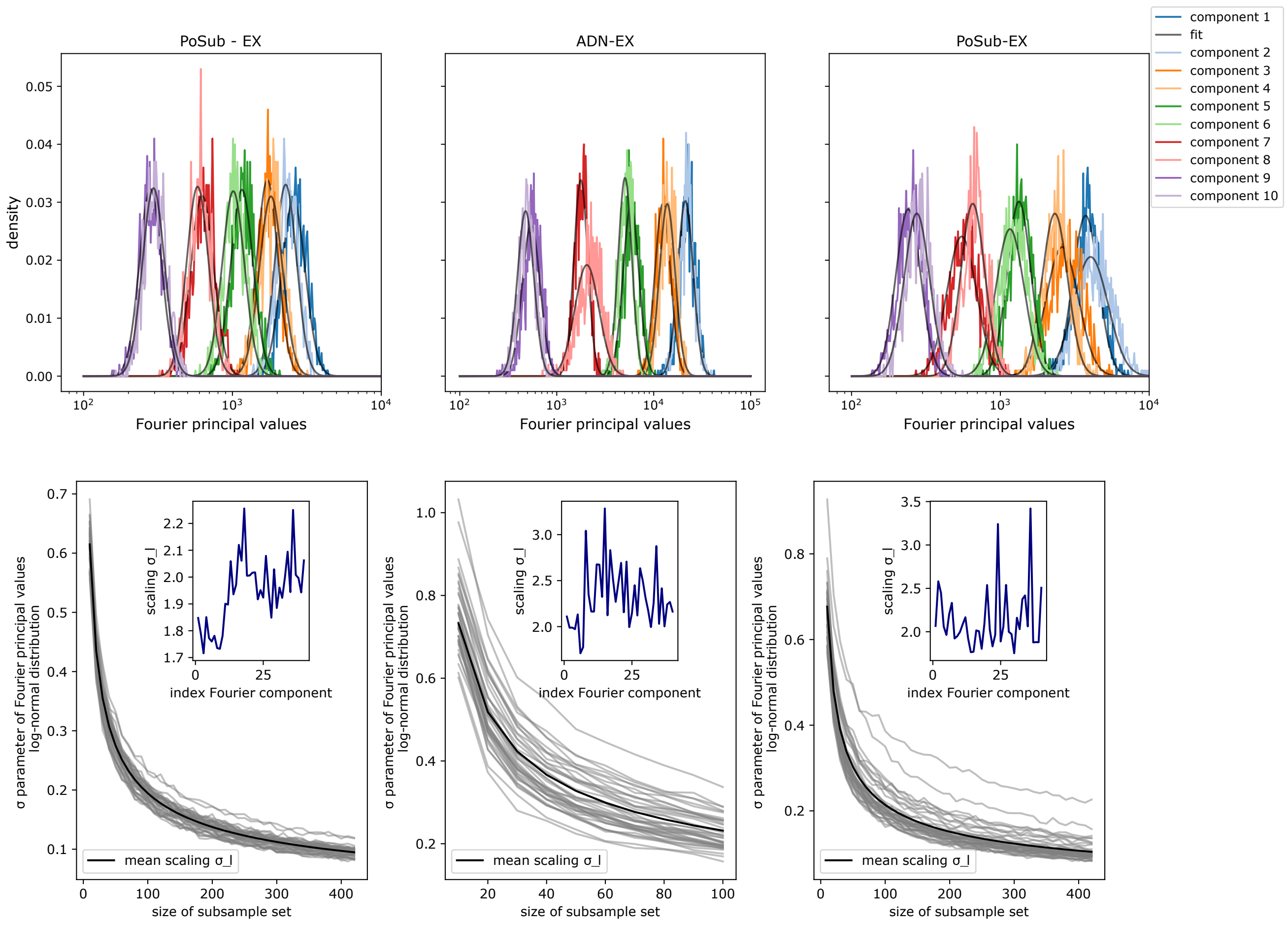


**Supplementary Figure 3.** Fitting the variance of Fourier principal values due to experimental sampling. Top: we show the log-normal fit of the Fourier principal values to a bootstrapped estimate of each Fourier principal value distribution. Bottom: evolution of the log-normal variance as a function of the subsample set size used in the bootstraping procedure. The fit is made on the HD dataset

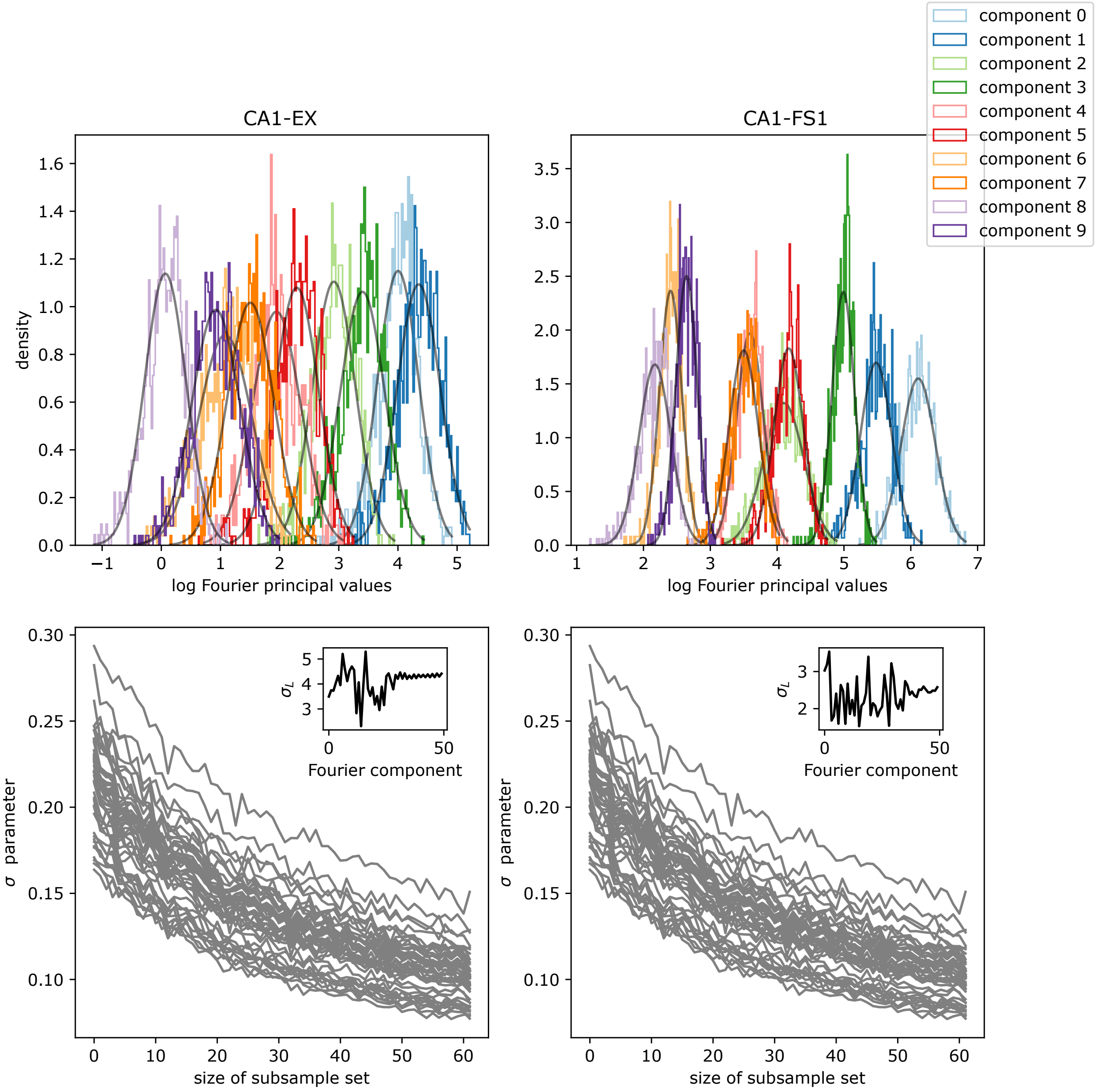


**Supplementary Figure 4.** CA1 dataset: Fitting the variance of Fourier principal values due to experimental sampling. Same as for the HD dataset.

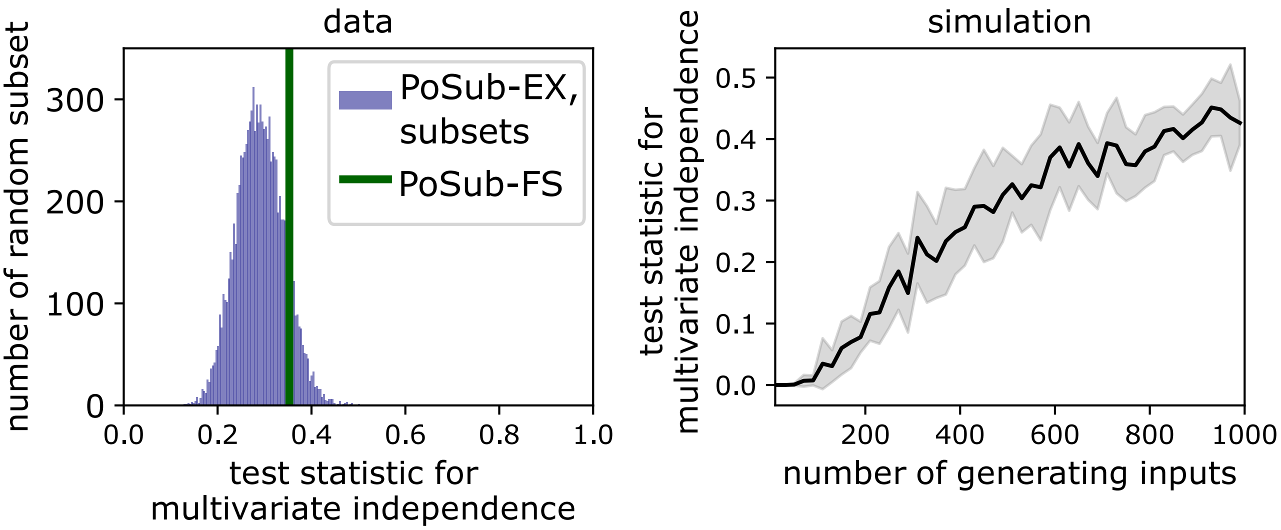


**Supplementary Figure 5.** Failure of the likelihood ratio test. Left, distribution of test statistic for multivariate independence when subsampling a random subset of pyramidal excitatory cells (blue). The green line is the test-statistic for the FS-cells tuning curves. Right, perfect random linear model where our theory applies exactly, generate test statistic that are in the range of the one computed from the data.

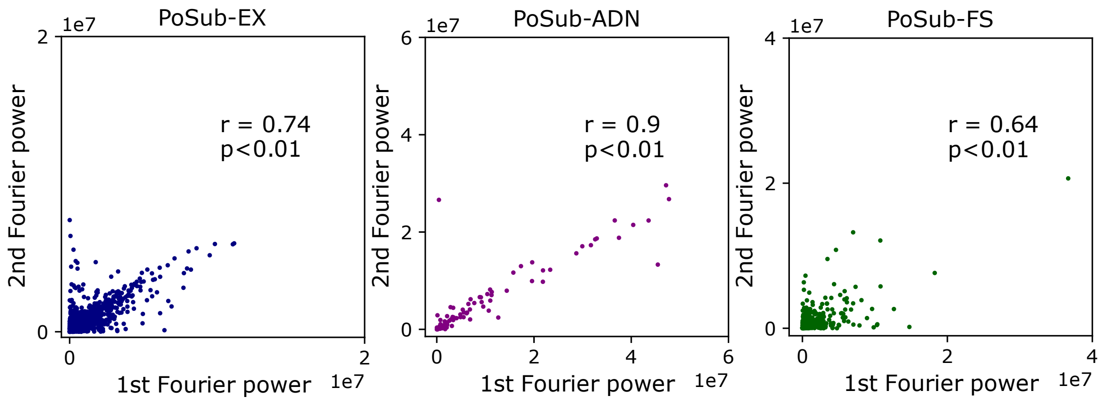


**Supplementary Figure 6.** Fourier power are not independent, even for FS cells. We compute the correlation between 1st and 2nd Fourier power for the three populations: pyramidal cells in Posub or AD and FS cells in Posub.

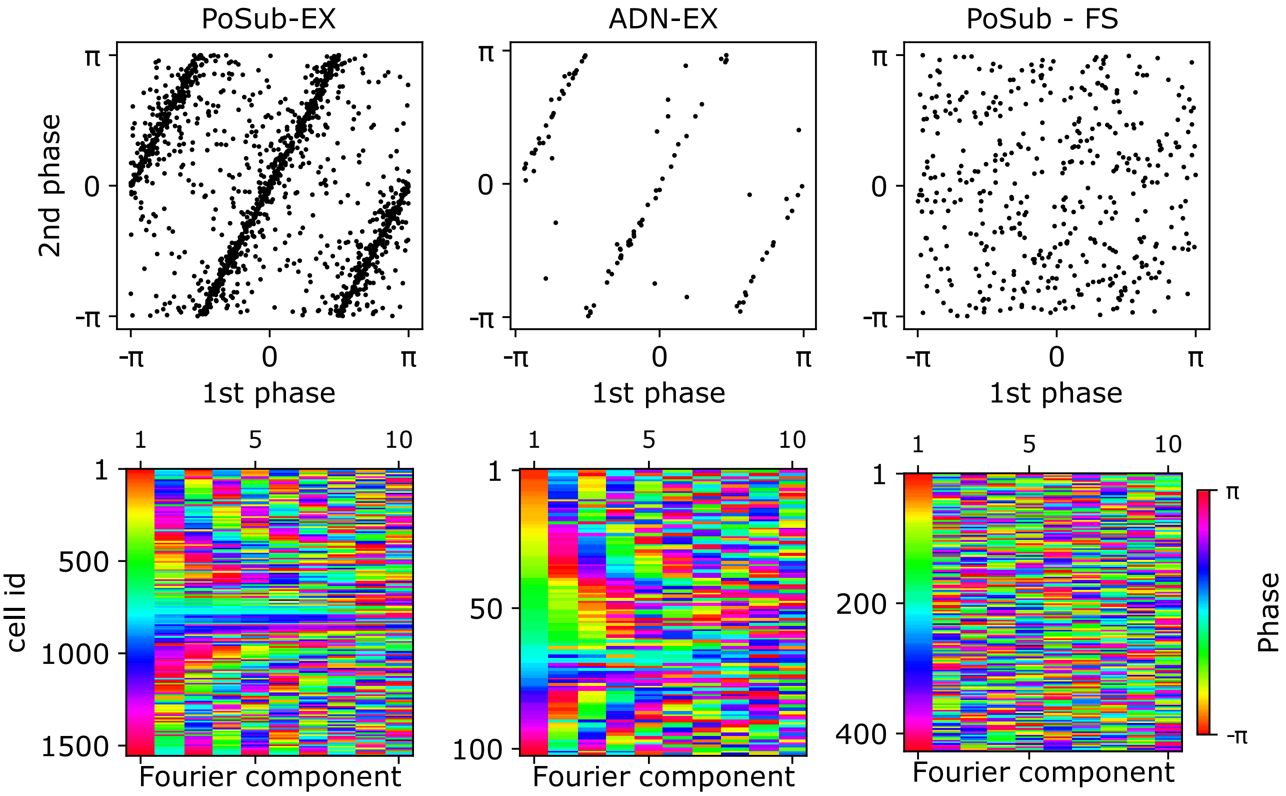


**Supplementary Figure 7.** Fourier phases are independent in FS cells but not HD cells. Top: scatter plor of the phase for Fourier component 1 and 2 in the three populations. Bottom: we sort the cells by the phase of their first Fourier component. The analytical link between the phase of the i-th Fourier component and the first Fourier component can be observed for excitatory cells but disappear for FS cells showing the independence of their phases.

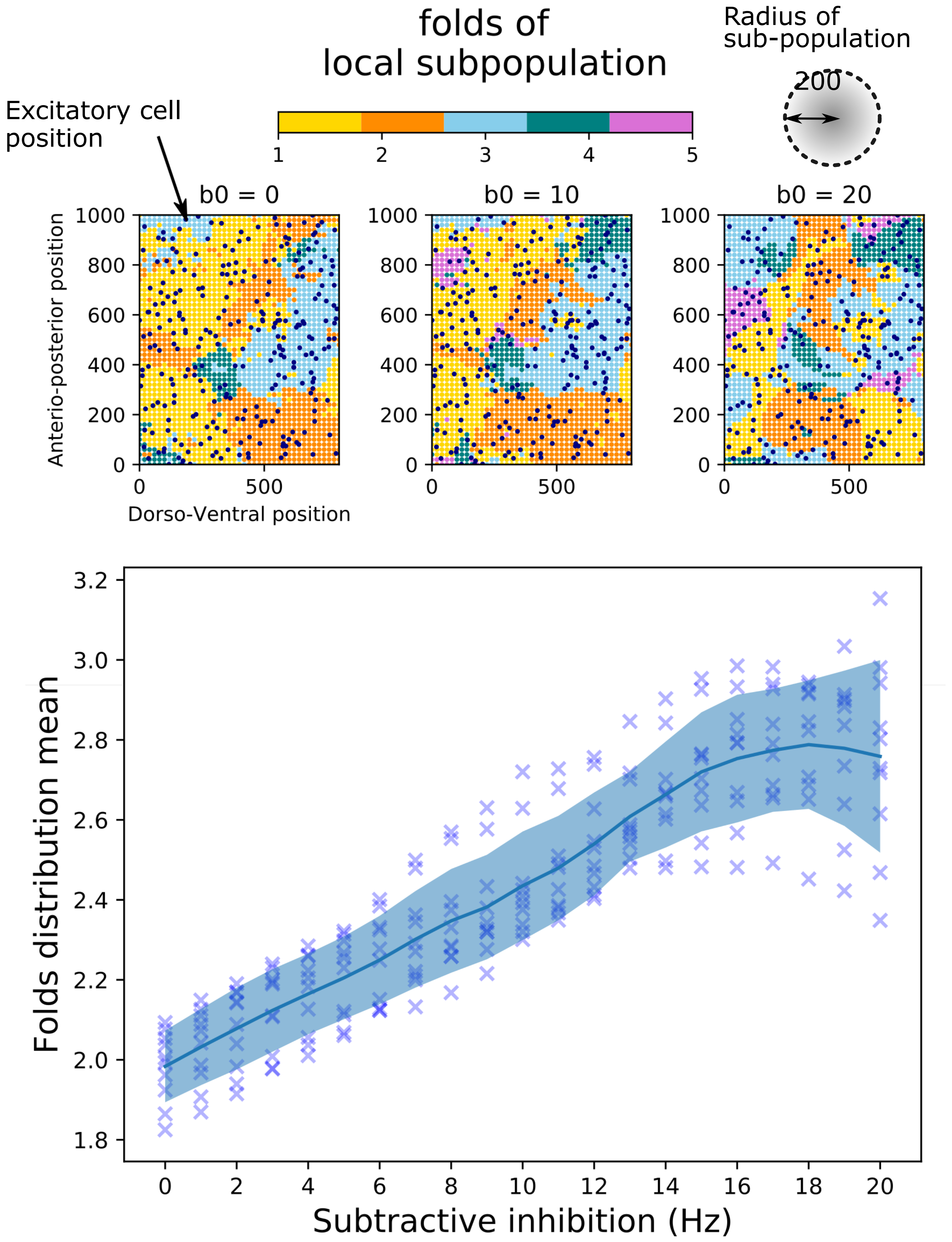


**Supplementary Figure 8.** The random linear model effect on a simulated cortical 2d sheet. We compute sub-population tuning curves by summing the tuning curves of excitatory neurons (relu non-linearity) in local sub-population of 200 $\mu m$. The mean number of folds across sub-population is plotted as a function of the global subtracting inhibition.
